# MetaboliteChat: A Unified Multimodal Large Language Model for Interactive Metabolite Analysis and Functional Insights

**DOI:** 10.1101/2025.11.07.687008

**Authors:** Zhenhao Guo, Dingcheng Duan, Youwei Liang, Atharvraj Patil, Pengtao Xie

**Affiliations:** Department of Computer Science, New York University, New York, NY 10012, USA; Department of Electrical and Computer Engineering, University of California San Diego, La Jolla, CA 92093, USA

## Abstract

Accurately characterizing the mechanisms and properties of metabolite molecules is essential for advancing metabolomics and systems biology. Yet most existing approaches are narrow, task-specific models that do not transfer across tasks and cannot express the richer, multimodal biological context of a metabolite in natural language. To address these challenges, we introduce MetaboliteChat, a multimodal ChatGPT-like large language model (LLM) that provides a unified and interactive framework for metabolite analysis. MetaboliteChat integrates molecular-graph and image reasoning with natural language understanding to generate comprehensive, free-form predictions about metabolite mechanisms and properties. The MetaboliteChat architecture consists of a graph neural network (GNN), a convolutional neural network (CNN), a large language model (LLM), and adapters, all trained end to end. This unified, multimodal design enables interactive reasoning over unseen metabolites, allowing the model to integrate structural and contextual cues and support discovery and translational insights across diverse biological systems.

## Introduction

Metabolomics holds substantial promise for biomedicine. Metabolites are closely associated with diverse human phenotypes, including health status, disease progression, aging, and therapeutic response (1–3). The ability to identify and characterize these molecules offers critical opportunities for developing new clinical strategies, drug monitoring techniques, and insights into human physiology (4–6). With recent advances in mass spectrometry, researchers can now generate high-resolution and high-sensitivity metabolomics data on a large scale, significantly expanding the potential for metabolic research and discovery (7, 8).

Despite this potential, conventional metabolite analysis is labor-intensive, costly, and dependent on extensive validation and manual curation. It also demands deep expertise in chemistry and biology to interpret structures and functions across a vast chemical space (9–12). At the model level, deep learning systems have emerged in recent years, but most remain task-specific and exhibit poor transferability (13, 14). For example, MetaboListem and TABoLiSTM target only named-entity recognition, lacking structural reasoning, knowledge-based inference, and downstream transfer (15). Moreover, architectures often require careful tailoring to each modality or objective, hindering practical use (16). On the data side, metabolomics is fundamentally multimodal: the same metabolite can be described through complementary views and modalities (17, 18). A growing body of work shows that integrating these modalities yields deeper biological insight than treating any single modality in isolation (19, 20). Nonetheless, most computational pipelines still operate on narrowly defined inputs and favor linear models such as PLS-DA, because such linear models are considered transparent and easy to interpret (21–23). This comes at the cost of missing nonlinear, cross-modal relationships, so biologically relevant complexity is often not captured.

Taken together, task-specific modeling, modality-specific architectural complexity, and the absence of multimodal reasoning have so far hindered the emergence of a unified, generalizable framework for metabolomics. To address them, we present MetaboliteChat, a ChatGPT-like system that enables free-text, multi-turn dialogues for exploring and reasoning about metabolites across multiple modalities. The interface supports structural interpretation, biological context, and functional inference, enabling users to ask open-ended questions about individual compounds, and obtain insights.

MetaboliteChat applies a multimodal framework in metabolomics by integrating graph-based and image-based representations with a language-model interface. A graph neural network (GNN) (24) and a convolutional neural network (CNN) (25) encode molecular structure and visual features; these embeddings condition an LLM (26, 27) that generates context-aware, free-form responses. This multimodal design improves interpretability and predictive utility by combining complementary molecular attributes and contextualizing their relationships. By aligning molecular structures, image representations, and text prompts in a unified, dialog-centric model, MetaboliteChat generalizes to previously unseen metabolites, producing biologically grounded, high-quality responses and outperforming general-purpose foundation models.

## Results

### MetaboliteChat Overview

MetaboliteChat is a multimodal LLM that generates textual predictions about metabolites based on their SMILES (Simplified Molecular Input Line Entry System) representation and user-provided prompts, as depicted in Figure 1. When given a metabolite’s SMILES string and a query like “Please give me some information about this metabolite.” the system can produce detailed responses such as “Convallatoxoloside is isolated from Convallaria majalis. Convallaria majalis is banned…” This capability makes MetaboliteChat a powerful tool for chemical and pharmaceutical research.

**Fig. 1.**
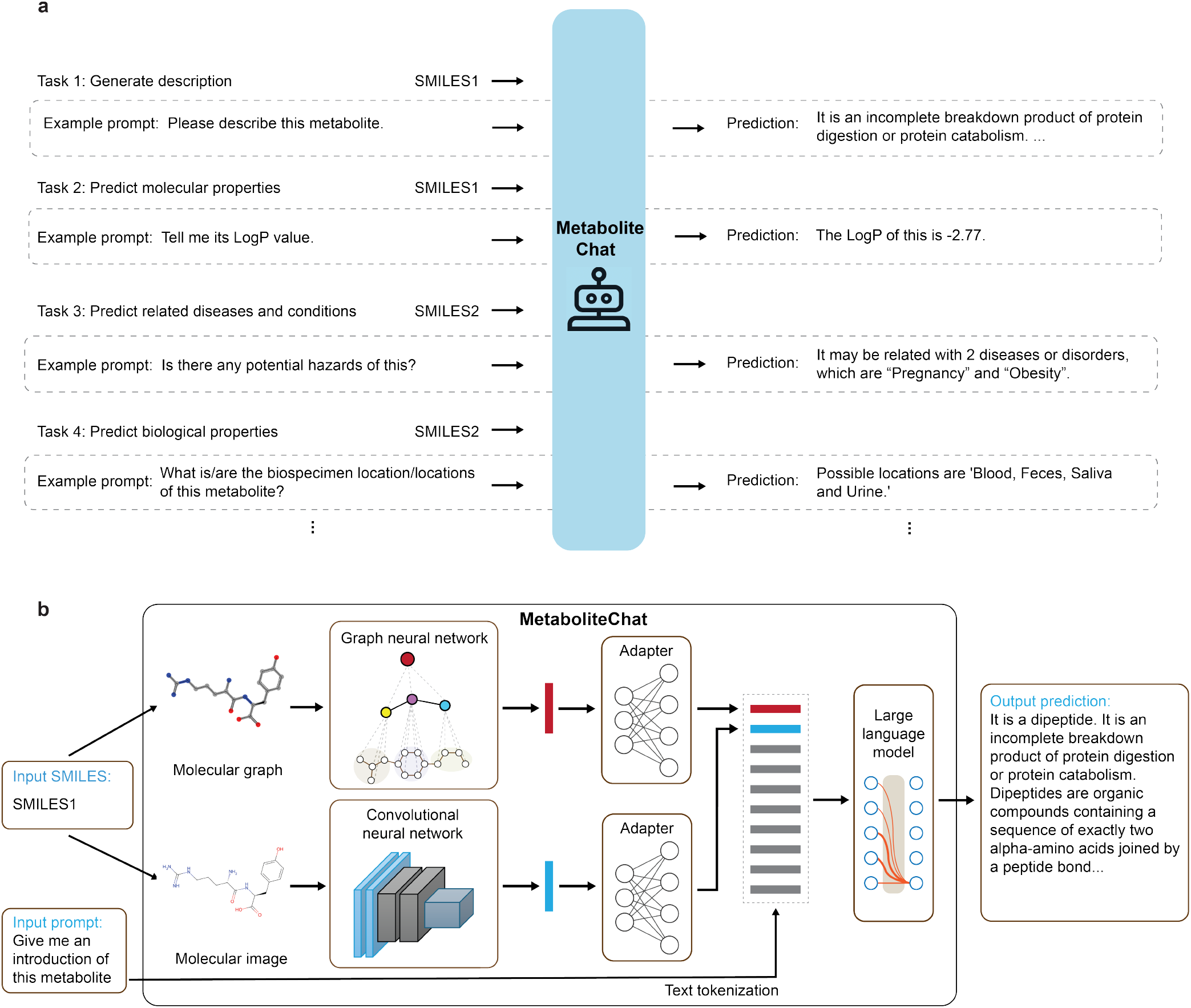
MetaboliteChat is a multimodal LLM for predicting metabolite attributes in free-form text or discrete categories. (a) The system supports diverse prediction tasks through natural language prompts, enabling flexible, task-specific queries without changing model parameters. (b) Model architecture: SMILES input is processed into molecular graph and image representations via a graph neural network and a convolutional neural network. Two adapters convert these representations into a format compatible with the large language model, which generates predictions based on the molecular features and the prompt. (c) Example SMILES inputs: smiles1: N[C@@H](CCCNC(N)=N)C(=O)N[C@@H](CC1=CC=C(O)C=C1)C(O)=O ssmiles2: CCCCCCC\C =C/CCCCCCCCCC(=O)O[C@](H)(COC(=O)CCCC\C =C/C\C =C/C\C =C/CCCCC)COP([O-])(=O)OCC[N+](C)(C)C

At a high level, MetaboliteChat links molecular structure to language by combining neural encoders that extract rich representations of a molecule, and a large language model that produces contextualized answers conditioned on both the molecule and the user’s prompt. As shown in Figure 1, a single SMILES string is processed along two complementary paths: one path converts the SMILES into a molecular graph, and the other into a rendered molecular image. These two views capture different but biologically important aspects of the metabolite.

For molecular encoding, the graph path uses a pretrained GNN trained with self-supervised learning on two million unlabeled molecules from ZINC15 (28, 29). The GNN encodes nodes and edges into feature vectors, aggregates neighborhood information, and yields a graph-level embedding that captures structural information. In parallel, the image path uses the pretrained ImageMol CNN (30), trained on images of ten million bioactive molecules from PubChem (31). The CNN extracts hierarchical visual features via 2D convolutions and pools them into a compact image representation. To integrate these molecular representations with language, multilayer perceptron (MLP) adapters map the graph and image embeddings into a unified molecule token compatible with the LLM, while the user’s text prompt is tokenized into language embeddings. The final processing stage employs Vicuna-13B (32), a powerful large language model, which processes the combined input through autoregressive decoding (33). This allows MetaboliteChat to generate coherent responses that integrate multimodal molecular information with the user’s intent, enabling meaningful insights through natural language interaction.

To train MetaboliteChat, we curated data from the Human Metabolome Database (HMDB) (34). We obtained 217,460 unique metabolites and constructed 351,138 (metabolite, prompt, answer) triplets based on the available informative attributes from HMDB. For evaluation, the 217,460 metabolites were split into training (70%), validation (20%), and test (10%) sets, using the former for optimization and the latter two to assess predictive accuracy and generalization.

### MetaboliteChat Free-form Predictions in Various Fields Outperform GPT-4o

To assess its capability, we conducted free-form prediction experiments comparing MetaboliteChat with GPT-4o using the test set. Starting with the corresponding molecule’s SMILES and a simple prompt like “Please describe this metabolite” or “Give me an introduction of this metabolite.”, MetaboliteChat is able to generate free-form answers to provide basic information about the metabolite (such as its structure, function and category). The evaluation metrics include: BLEU (35), ROUGE (36), and Semantic Similarity scores. BLEU scores assess n-gram overlap between model-generated outputs and ground-truth annotations and reflect stringent requirements for phrase-level fluency and contextual coherence. ROUGE-1 measures unigram recall, complementing BLEU by focusing on how much of the reference content is covered by the generation. ROUGE-L evaluates the longest common subsequence, capturing both content fidelity and structural alignment.

Across all metrics, MetaboliteChat consistently outperforms GPT-4o. For BLEU-1/2/3/4, it achieves 0.3576, 0.2992, 0.2654, and 0.2277, respectively, compared to 0.0418, 0.0171, 0.0085, and 0.0051 for GPT-4o, as shown in Figure 2. The boxplots make the pattern explicit: for Semantic Similarity, ROUGE-1, and ROUGE-L, MetaboliteChat’s median scores are markedly higher, its lower quartile sits above GPT-4o’s upper quartile, and its upper whiskers approach 1.0, whereas GPT-4o’s distribution is concentrated near zero, as demonstrated in Figure 3. Together, these shifts indicate consistent structural fidelity and semantic alignment, and they demonstrate that MetaboliteChat delivers robust performance on unseen metabolites, outperforming GPT-4o across all metrics.

**Fig. 2.**
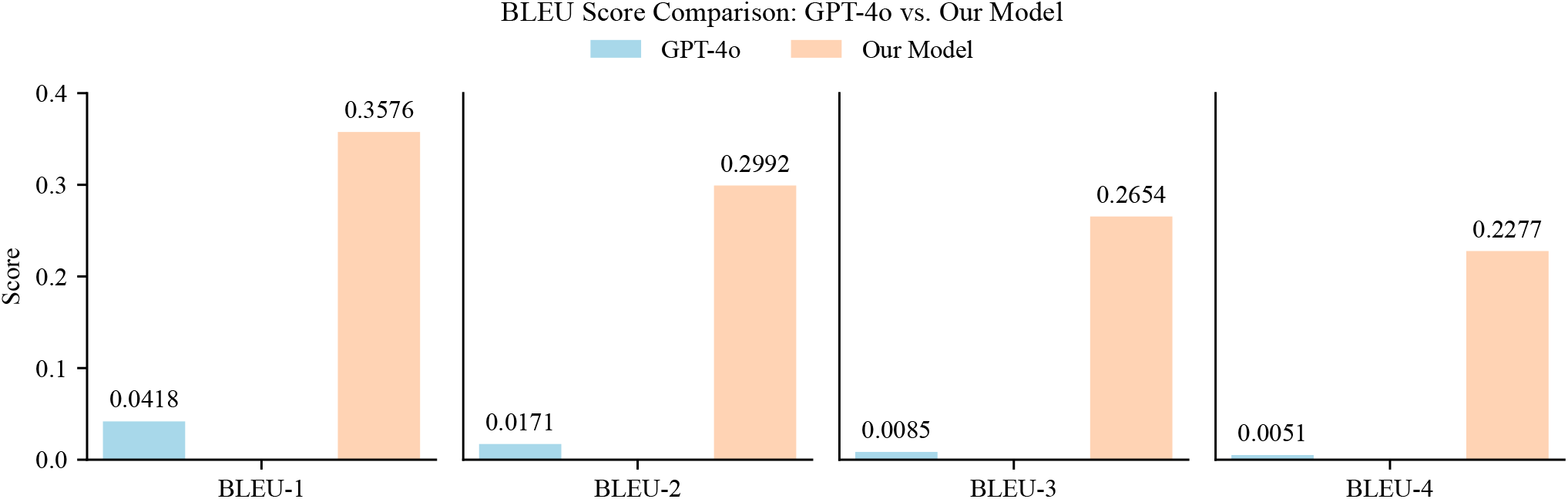
MetaboliteChat substantially outperforms GPT-4o across BLEU metrics. Bar charts compare GPT-4o and MetaboliteChat on BLEU-1, BLEU-2, BLEU-3, and BLEU-4. MetaboliteChat attains scores of 0.3576, 0.2992, 0.2654, and 0.2277, respectively, versus 0.0418, 0.0171, 0.0085, and 0.0051 for GPT-4o (Table 1).

**Fig. 3.**
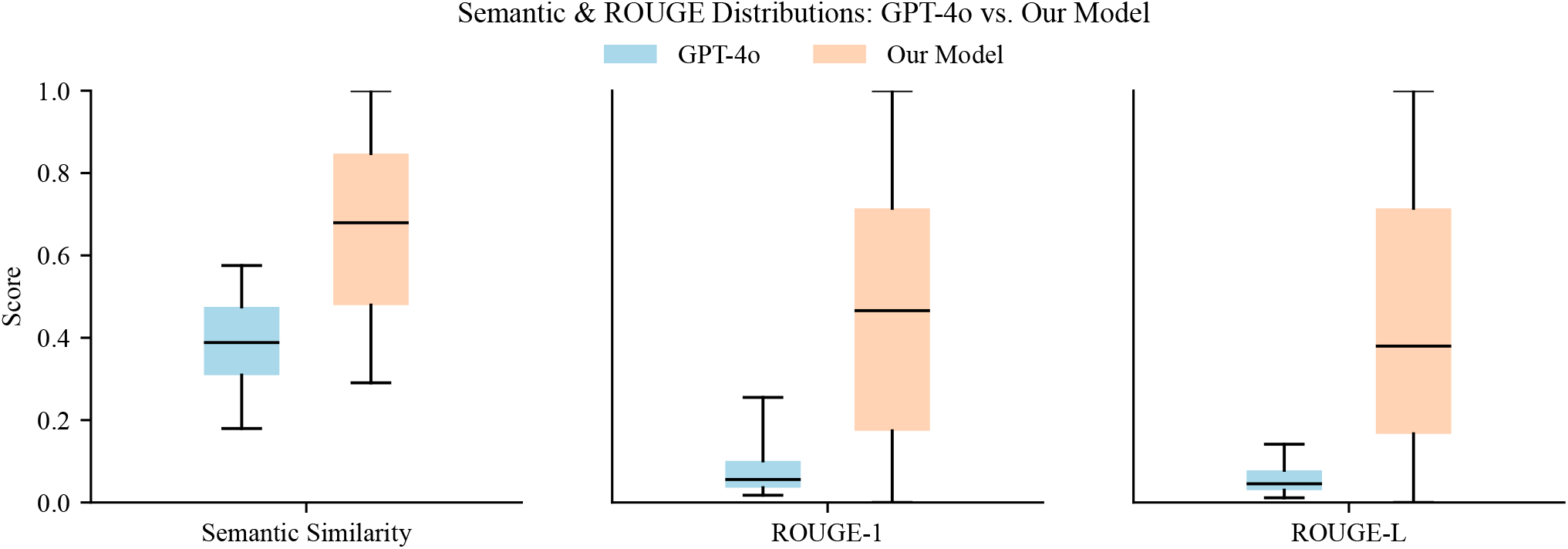
MetaboliteChat achieves higher semantic and ROUGE than GPT-4o. Boxplots compare GPT-4o and MetaboliteChat on Semantic Similarity, ROUGE-1, and ROUGE-L. Across all metrics, MetaboliteChat’s distributions shift substantially upward: its medians are consistently higher, its lower quartiles exceed GPT-4o’s upper quartiles, and its upper whiskers approach 1.0. In contrast, GPT-4o’s distributions are concentrated near zero, indicating limited semantic alignment and structural preservation (Table 2).

### MetaboliteChat Enables Interactive and Iterative Predictions of Metabolite Functions

MetaboliteChat facilitates interactive dialogues between users and the system. This interactive mechanism enhances both the transparency and reliability of the model. After obtaining initial predictions from MetaboliteChat, users can prompt with more detailed and specific questions to further refine and expand these predictions. Figure 4 shows example exchanges that highlight its versatility: When asked about the bio-specimen locations of a metabolite, the system provides the correct context; when prompted for its LogP (octanol–water partition coefficient), it returns the accurate value; and when broader information is requested, it generates a comprehensive description covering the metabolite’s chemical classification, biological functions, and metabolic pathways. This example highlights MetaboliteChat’s ability to flexibly shift between concise factual answers and broader explanatory descriptions. Unlike previous methods limited to single-shot predictions, MetaboliteChat enables iterative exploration of metabolite functions. Through sustained dialogues, users can progressively refine questions and obtain both accurate and comprehensive insights into metabolite behaviors and biological mechanisms, as demonstrated in Figure 5.

**Table 1.** Mean evaluation scores comparing our model and GPT-4o on BLEU metrics.

| Model | BLEU-1 | BLEU-2 | BLEU-3 | BLEU-4 |
| --- | --- | --- | --- | --- |
| Our Model | 0.3576 | 0.2992 | 0.2654 | 0.2277 |
| GPT-4o | 0.0418 | 0.0171 | 0.0085 | 0.0051 |

**Table 2.** Mean values for the metrics shown in Figure 3.

| Model | Semantic | ROUGE-1 | ROUGE-L |
| --- | --- | --- | --- |
| Our Model | 0.6547 | 0.4498 | 0.4234 |
| GPT-4o | 0.3869 | 0.0873 | 0.0597 |

**Fig. 4.**
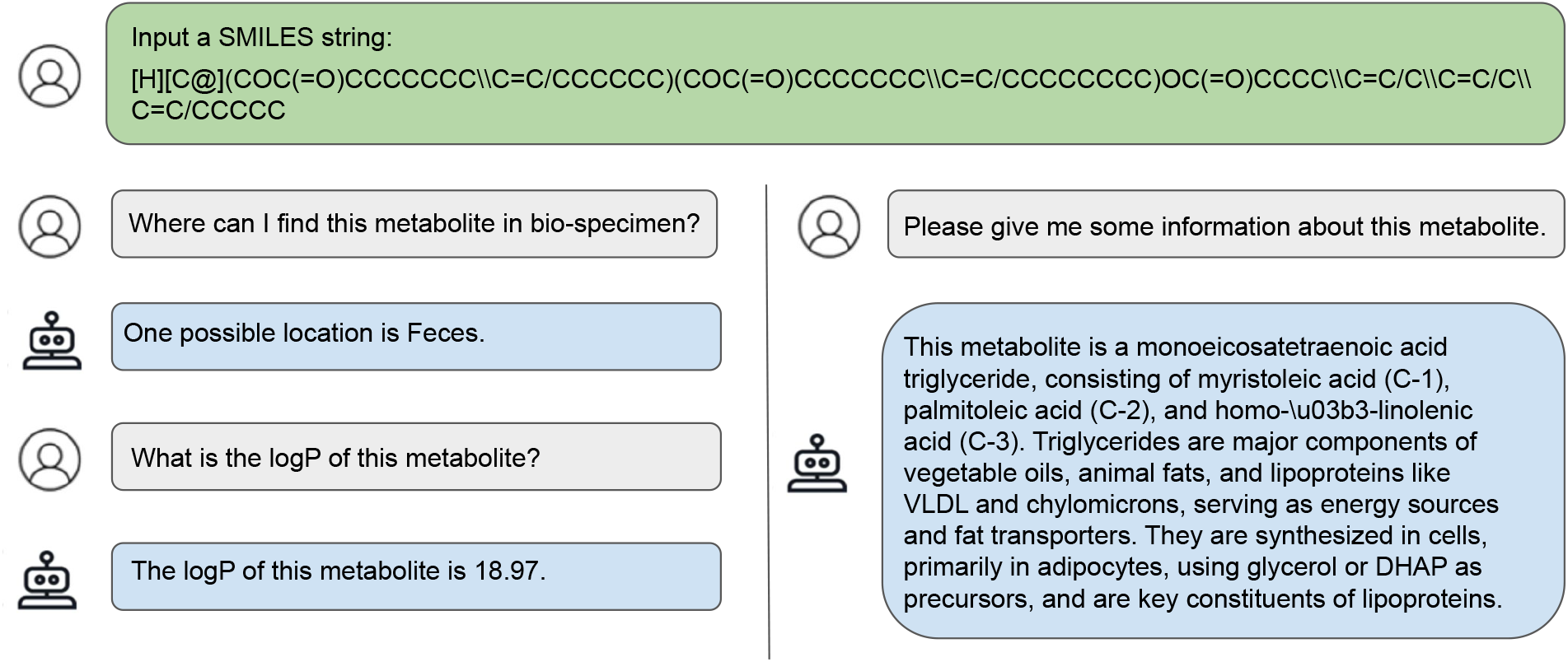
An exemplar multi-turn dialogue between MetaboliteChat and a user regarding the same metabolite.

**Fig. 5.**
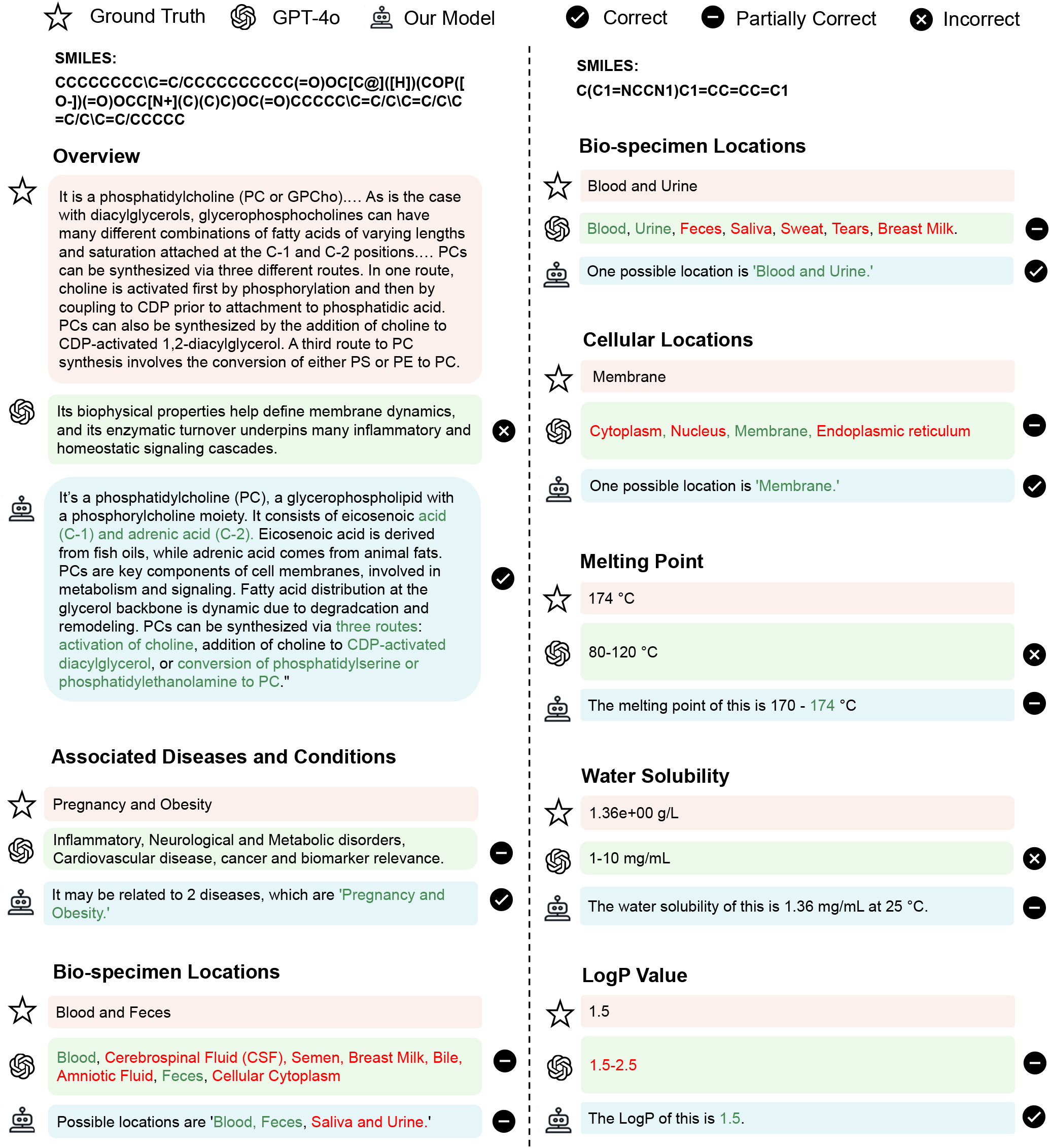
Examples comparing GPT-4o and *MetaboliteChat* in free-form predictions, showing our model’s superior performance and accuracy. The figure illustrates cases where MetaboliteChat’s generated descriptions align more closely with ground-truth annotations across various biomedical attributes, demonstrating improvements in both correctness and completeness.

## Discussion

The key contributions of this paper are threefold. First, MetaboliteChat is a multimodal LLM that ingests molecular graphs and images alongside text, enabling it to perceive structural connectivity, stereochemistry, and visual cues. By incorporating these inputs rather than relying solely on textual encodings such as SMILES, the system preserves richer structural information. This fusion yields mechanism-aware interpretations and improves predictive utility, while retaining the generative and reasoning strengths of an LLM. Together, these properties enable grounded answers with multimodality perspectives rather than simple pattern matching. Second, beyond multimodal fusion, our contribution centers on interactive and explanatory workflows. It answers open-ended queries by identifying candidate metabolites, explaining its reasoning, suggesting follow-ups, and adapting responses. In practice, this versatility benefits multiple settings. For researchers, MetaboliteChat can interpret proposed structures or images, suggest plausible compound identities, and generate concise, task-oriented summaries. For clinicians, it can generate accessible biology overviews and flag potential pathway or drug associations. Its ability to generate explicit, human-readable rationales makes it accessible to both experimental researchers and clinical end users. Third, MetaboliteChat demonstrates strong generalization to unseen metabolites. It produces biologically grounded, structurally consistent explanations for metabolites it has never seen. This behavior is relevant in metabolomics settings where many metabolites are only partially annotated, lack established reference entries, or originate from rare or experimentally derived compounds.

While multimodal fusion and multi-turn dialogue enhance reliability, fine-tuning the model on HMDB provides valuable domain knowledge. However, it can also introduce dataset-specific biases, yielding confident yet incorrect predictions on unfamiliar metabolites. Mitigations include coupling the model with deterministic cheminformatics tools, using spectral libraries, and reporting uncertainty explicitly. Adding modalities would further improve reliability.

## Methods

### Dataset Preprocessing

To train MetaboliteChat, we collected data from HMDB, with the distribution shown in Figure 6. After collection and cleaning, we obtained 217,460 unique metabolites stored in a structured format for training. For each metabolite, we extracted up to eight properties when available: (1) SMILES, the molecular structure representation; (2) Cellular Locations, where in the cell the metabolite is found; (3) Biospecimen Locations, the bio-specimens (e.g., blood, urine) in which the metabolite is present; (4) Tissue Locations, the human tissues containing the metabolite; (5) Description, a textual summary of properties and biological significance; (6) Diseases, disease associations, including biomarker roles;(7) Experimental Molecular Properties, such as solubility, pKa, and other physicochemical characteristics; and (8) Experimental Chromatographic Properties, detailing chromatographic behavior. These attributes are not uniformly populated across metabolites, so only available fields were used to build the triplets for training and evaluation.

**Fig. 6.**
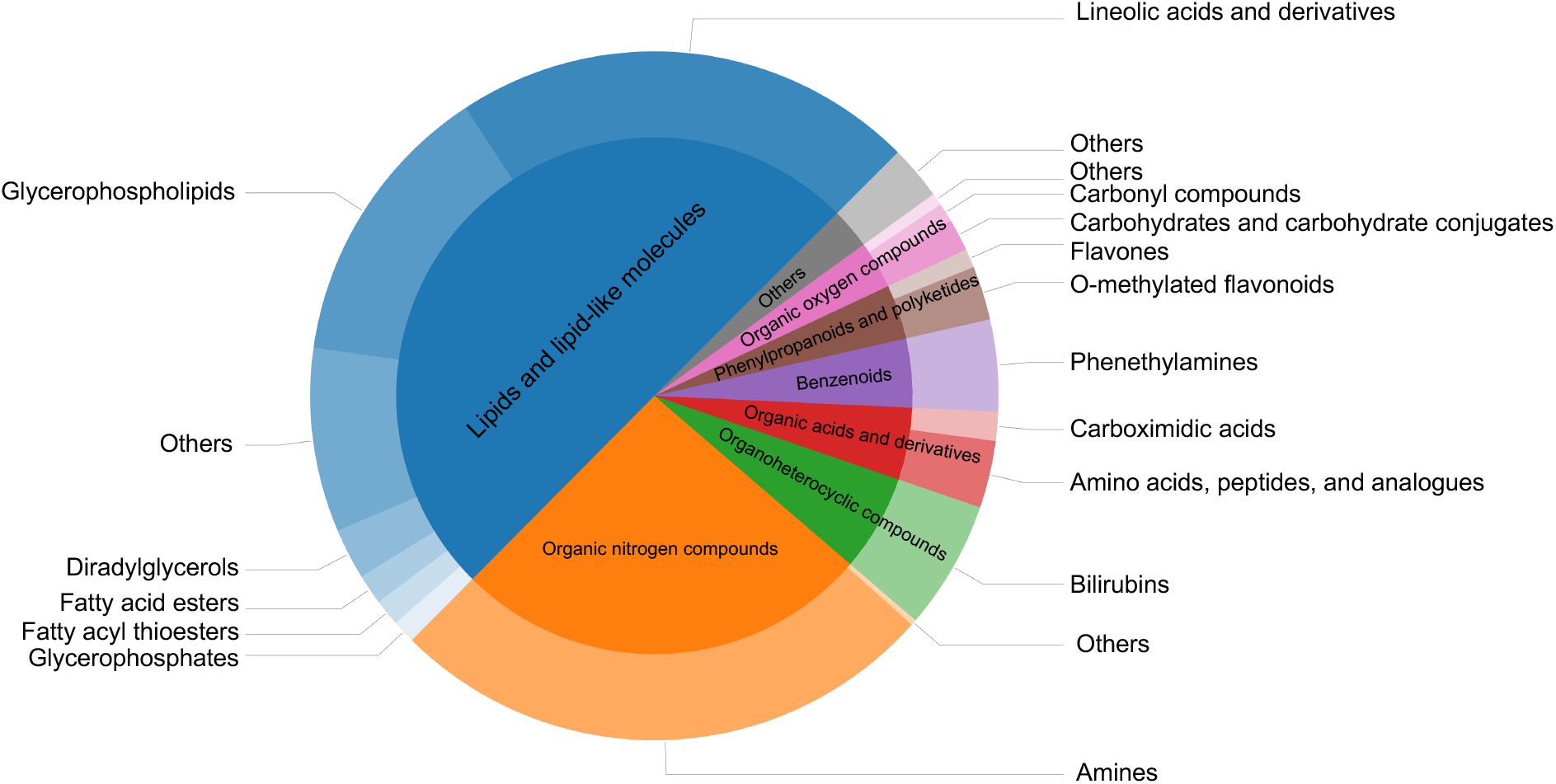
The figure represents the distribution of categories in the dataset. The inner disk represents the superclasses, while the outer ring shows the corresponding subclasses, highlighting the diversity of metabolic space.

Using these metabolites and their annotated attributes, we curated the training data for MetaboliteChat. For each attribute of a metabolite, we created a triplet consisting of the metabolite’s SMILES representation, a textual prompt querying the value of the attribute, and the corresponding ground truth value. Each attribute type had its own tailored prompt. For example, for the metabolite with the SMILES string “NC12CC3CC(CC(C3)C1)C2,” the prompt “Please describe this metabolite.” would be paired with the value “An antiviral that is used in the prophylactic or symptomatic treatment of influenza A. It is also used as an antiparkinsonian agent, to treat extrapyramidal reactions, and for postherpetic neuralgia. The mechanisms of its effects in movement disorders are not well understood but probably reflect an increase in synthesis and release of dopamine, with perhaps some inhibition of dopamine uptake.” Other properties like cellular location, tissue location, and disease associations were queried using tailored classification prompts, such as “Where can I find this metabolite in bio-specimen?” or “What are the associated disorders and diseases with this metabolite?” The corresponding values were given as a list then added to form complete training examples.

### MetaboliteChat Model

MetaboliteChat is a multimodal model that fuses three modalities via a GNN, a CNN, and the LLM. For each molecule, RDKit^1^ processes the SMILES string to generate both a molecular graph and a molecular image.

In the molecular graph, nodes are atoms and edges are chemical bonds. Each atom is represented by its atom type (120 categories, including “Unknown”) and chirality (tetrahedral clockwise, tetrahedral counter-clockwise, unrecognized, other). Each bond is represented by type (single, double, triple, aromatic) and direction (none, upward, downward). All categories are embedded as learnable vectors. The graph is fed to a pretrained graph neural network. At each layer, the GNN updates node representations by aggregating neighboring nodes and edges; after *K* layers, information has propagated along *K*-hop paths. The graph embedding is the mean of node representations after *K* layers. Our GNN has five layers (approximately 0.5 million parameters); node and edge embeddings are 300-dimensional. The GNN was pretrained with context prediction (37) on two million unlabeled ZINC15 molecules. For the molecular image, we use a pretrained ResNet-18 (ImageMol) (38) with 18 convolutional layers and 11 million parameters to extract a 512-dimensional embedding.

After extracting representation vectors from the molecule using the GNN and CNN, we apply two separate MLPs, referred to as the adapters, to transform these molecular representations into a format compatible with the LLM, aligning them to the language space (39). The LLM typically uses Transformer decoders (40) to model natural language as a sequence of tokens, where each token is represented as a vector (41). In MetaboliteChat, the transformed molecular representations are treated as tokens and appended to the language token sequence, which was converted from the input prompt. This combined sequence is then passed into the LLM, which uses multi-head self-attention mechanisms to generate new tokens. These generated tokens form the final prediction. MetaboliteChat uses Vicuna-13B as the LLM, which contains 13 billion parameters. It was fine-tuned from Llama-13B (42) on a dataset of 70K user-shared dialogues collected from ShareGPT.com (containing conversations between human users and ChatGPT). The adapters responsible for converting molecular graph representations into LLM tokens consist of two layers, with an input dimension of 300, a hidden dimension of 5120, and an output dimension of 5120, totaling 28M parameters. Similarly, the adapters for converting molecular image representations include two layers, with an input dimension of 512, a hidden dimension of 5120, and an output dimension of 5120, amounting to 29M parameters. Both MLPs use the GELU activation function (43) in the hidden layer.

Building upon the architectural components described above, we formulate the training objective used to optimize MetaboliteChat. For a target answer *T* that has *L* text tokens, MetaboliteChat computes the probability of generating *T* as follows:

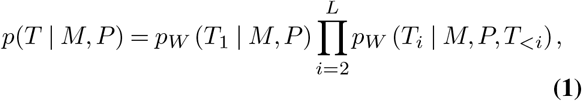

where *M* represents the input molecule and *P* is the input prompt. We denote the *i*-th token as *T*_*i*_ and all preceding tokens as *T*_*<i*_. Model parameters are denoted by *W*. The generated token sequence is compared to the ground truth tokens to compute the negative log likelihood (NLL). The parameters *W* are optimized by minimizing the sum of NLL over all training data.

### Model Training

We trained MetaboliteChat using the AdamW optimizer with 20 epochs, employing default parameters *β*_1_ = 0.9, *β*_2_ = 0.999 and a weight decay of 0.03. In the first 10,000 steps, we applied a linear warmup strategy, gradually increasing the learning rate from 10^−6^ to 3 *×* 10^−5^, which was then reduced to 10^−6^ using a cosine decay scheduler over the remaining training process. Considering the token length limit, we set the batch size to 4, and training was performed on an NVIDIA A100 GPU with 80 GB memory.

### Evaluation Metrics

We evaluated the quality of the generated metabolite descriptions using three widely used metrics in natural language generation tasks: BLEU, ROUGE, and Semantic Similarity scores.

#### BLEU (Bilingual Evaluation Understudy)

score measures the overlap between predicted and reference texts using modified n-gram precision, along with a brevity penalty to penalize excessively short outputs. It is computed as:

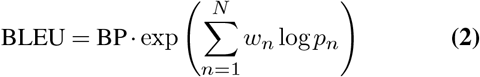

where *p*_*n*_ is the modified precision for n-grams of length *n*, and *w*_*n*_ is the corresponding weight (typically uniform). The brevity penalty (BP) discourages short predictions and is defined as:

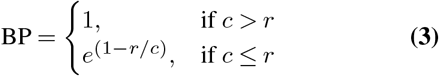

where *c* is the length of the predicted text and *r* is the length of the reference text.

#### ROUGE (Recall-Oriented Understudy for Gisting Evaluation)

scores complement BLEU by emphasizing recall and content coverage. ROUGE-N measures n-gram recall and is computed as:

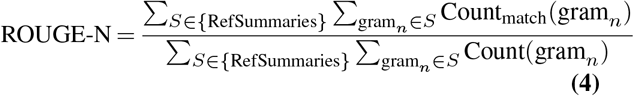

where Count_match_(gram_*n*_) is the maximum number of ngrams co-occurring in a candidate summary and reference summary, and Count(gram_*n*_) is the number of n-grams in the reference summary. ROUGE-L evaluates the longest common subsequence (LCS) between generated and reference texts:

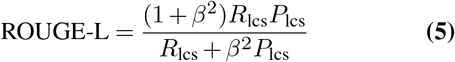

where *R*_lcs_ and *P*_lcs_ are LCS-based recall and precision respectively, and *β* controls the relative importance of recall versus precision.

#### Semantic Similarity

was calculated as the cosine similarity between the sentence embeddings of the ground-truth and model-predicted texts, with embeddings generated using the pretrained sentence transformer model All-MiniLM-L6-v2 (44). Let *SE* denote the sentence embedding model. For a ground-truth text *t*_*g*_ and a predicted text *t*_*p*_, their embeddings are *e*_*g*_ = *SE*(*t*_*g*_) and *e*_*p*_ = *SE*(*t*_*p*_), respectively. The similarity is defined as::

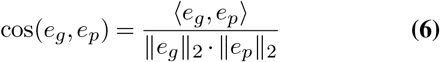

where ⟨*e*_*g*_, *e*_*p*_⟩ is the dot product and ∥ · ∥_2_ is the L2 norm. Higher values indicate greater semantic closeness in the embedding space.

### Experimental Details for the GPT-4o Baseline

The prompts used to query the model included both openended and fact-oriented questions. Depending on the input format, the model was queried either with metabolite names or with SMILES representations. When provided with a metabolite name, the prompt was: “Given the metabolite name [metabolite name], please give me some information about this metabolite.” When provided with the SMILES sequence, the prompt was: “Given the sequence of the SMILES: [a SMILES string such as CN1C=NC(C[C@H](N)C(O)=O)=C1], please give me some information about this metabolite.”

For fact-oriented tasks, direct queries were posed targeting specific physicochemical or biological properties of the metabolite. Examples include: “What is the water solubility of this metabolite?”, “What is the LogP of this metabolite?”, “Where is this metabolite in cell?”, “Where can I find this metabolite in bio-specimen?”, and “What is the melting point of this metabolite?”. Because certain queries correspond to classification tasks, we provided a list of possible candidates and instructed GPT-4o to select the most appropriate answer. Each of these property-related questions was phrased in a consistent format to ensure comparability across metabolites.

## Data Availability

All data used in this study are available at https://drive.google.com/drive/folders/1YYTQUJbYBVVoQ8B_P1ZYCUg8JuePQG_L?usp=sharing.

## Code Availability

The source code of this work is available at https://github.com/13501274828/MetaboliteChat

## Footnotes

1 https://www.rdkit.org

